## Supplementary material for "Integrated Molecular Characterisation of the MAPK Pathways in Human Cancers": Description of Supplementary Files (from another version of this preprint: version 2, 2020-05-04)

**Description of Additional Supplementary Files**

**Supplementary Data 1:** Supplementary data of cancer studies and mutations of the MAPK pathway genes. The spreadsheet contains the following results/datasets according to the sheet name. ***MAPK Pathway Genes;*** list of MAPK pathway genes that we curated from the literature including: the abbreviated name of the encoded protein, the official HUGO gene symbol, the signalling pathway module in which the gene participates and the class of MAPK pathway protein encoded by the genes. ***Cancer studies;*** list and description of 193 individual cancer studies from which our analyses are based. ***Specific MAPK Gene Mutations***; mutations that we found in each MAPK pathway gene across all the cancer studies. ***Cancer-MAPK Gene Mutations;*** the frequency of mutations that we found in each MAPK pathway gene involved for each cancer types. ***Cancer-Protein Type Mutations;*** mutations of each category of genes that encode a specific class of MAPK proteins across each cancer type. ***Cancer-MAPK pathways Mutations***; overall mutations frequencies across all the four MAPK pathway modules.

**Supplementary Data 2:** Clinical outcomes across various groups: The spreadsheet contains the following results/datasets according to the sheet name. ***Pathways - OS Pairwise Comp;*** Pairwise comparisons calculated using the Log-rank test ^1^ for the duration of overall survival periods between patients that had tumours with mutations in gene of, (1) only one of the four MAPK pathway modules, (2) in multiple MAPK pathway modules, (3) no mutations to the MAPK pathway genes (see Figure 2f). ***Proteins - OS Pairwise Comp;*** Pairwise comparisons calculated using the Log-rank test for the duration of overall survival periods between patients that had tumours with mutations in genes of, (1) only one class of MAPK encoding genes, (2) in multiple classes of MAPK encoding genes, (3) no mutations to any of these genes (see Figure 2g). ***Pathways - DFS Pairwise Comp;*** Pairwise comparisons calculated using the Log-rank for the duration of disease-free survival periods between patients that had tumour with mutations in gene of, (1) only one of the four MAPK pathway module, (2) in multiple MAPK pathway modules, (3) no mutations to any of the MAPK pathway genes (see Figure 2h). ***Proteins - DFS Pairwise Comp;*** Pairwise comparisons calculated using the Log-rank for the duration of disease-free survival periods between patients that had tumour with mutations in genes of, (1) only one class of MAPK encoding genes, (2) in multiple classes of MAPK encoding genes, (3) no mutations to any of these genes.

**Supplementary Data 3:** Achilles fitness screens across cancer types: The spreadsheet contains the following results/datasets according to the sheet name. ***MAPK vs MAPK Genes Dependence;*** mean difference comparison results between the pooled CRISPR-derived gene dependence scores of each cancer types for a particular MAPK signalling pathway module against the pooled CRISPR-derived gene dependence scores of the all MAPK pathway genes across all cancer types. ***MAPK vs Other Genes Dependence;*** mean difference comparison results between the pooled CRISPR-derived gene dependence scores of each cancer types for a particular MAPK signalling pathway module vs the pooled CRISPR-derived gene dependence scores of all genes that are not MAPK pathway genes. ***Gene Class- MAPK vs MAPK Dep;*** mean difference comparison results between the pooled CRISPR-derived gene dependence scores of each cancer types for genes that encode the classes of MAPK pathway protein versus the pooled CRISPR-derived gene dependence scores for all the MAPK pathway genes across all the 688 cell lines. ***Gene Class- MAPK vs Other Genes;*** mean difference comparison results between the pooled CRISPR-derived gene dependence scores of each cancer types for genes that encode the classes of MAPK pathway protein versus the pooled CRISPR-derived gene dependence scores for all other genes that are not MAPK pathway genes. ***The dependency of MAPK Genes;*** the number of cell lines whose fitness score fall below -0.5 after CRISPR gene knockout. ***Fitness-Expression Correlation;*** linear Pearson’s correlation between the CRISPR-derived gene dependence scores and gene’s self-mRNA for all proteins coding genes that have CRISPR fitness data. ***GO Bio Process CRISPR NEG corr;*** gene ontology biological processes enriched for gene set that shows a negative linear Pearson's correlation score < -0.30 between the CRISPR-derived gene dependence scores and gene’s self-mRNA. ***GO Bio Process CRISPR POS corr;*** gene ontology biological processes enriched for gene set that shows a positive linear Pearson's correlation score > 0.30 between the CRISPR-derived gene dependence scores and gene’s self-mRNA. ***CNA Alterations;*** the number of copy number variations that we found in each gene involved in the MAPK pathways across all the cancer types.

**Supplementary Data 4:** Dose-response of the cancer cell lines: The spreadsheet contains the following results/datasets according to the sheet name. ***MAPK Drug GDSC Dose Response;*** data on the dose-responses of the cancer cell lines to the 28 drugs that target the MAPK pathways. ***Between Cell line Dose Responses;*** mean difference comparison of the dose-responses to MAPK pathway inhibitors between the cancer cell lines that have a higher dependence on MAPK signalling and those with a lower dependence on MAPK signalling as defined using the CRISPR-derived gene dependence scores (see methods section). ***Between Cancer Drug Responses;*** mean difference comparison of the dose-responses to MAPK pathway inhibitors between the cancer types that have a higher dependence on MAPK signalling and those with a lower dependence on the MAPK signalling. ***CRISPR Drug Sensitive Assoc;*** mean dose-response comparison between cell lines that have a higher dependence on a particular MAPK pathway gene versus those that have a lower dependence on that MAPK pathway gene (see methods section). ***CRISPR Drug Sensitive Genes;*** genes that are associated with significantly increased sensitivity to various MAPK pathway inhibitors in the cell lines that show a higher dependence on the particular gene(s) for their fitness. ***CRISPR Drug Resistance Genes;*** genes that are associated with a significant resistance to various MAPK pathway inhibitors in the cell lines that show a higher dependence on the particular gene(s) for their fitness. ***Mutations Gene Drug Response;*** mean dose-response comparison between cell lines that have a mutation(s) in a particular MAPK pathway gene versus those that do not have a mutation(s) in that particular MAPK pathway genes. ***Mutations Drug Sensitive Genes;*** genes that are associated with significantly increased sensitivity to various MAPK pathway inhibitors in the cell lines that have mutations in the particular gene. ***Mutations Drug Resistant Genes;*** genes that are associated with significant resistance to various MAPK pathway inhibitors in the cell lines that have mutations in the particular gene.
