## Supplementary Information revised-1 for "Integrated Molecular Characterisation of the MAPK Pathways in Human Cancers" (from another version of this preprint: version 2, 2020-05-04)

### Supplemental Information Titles and Legends

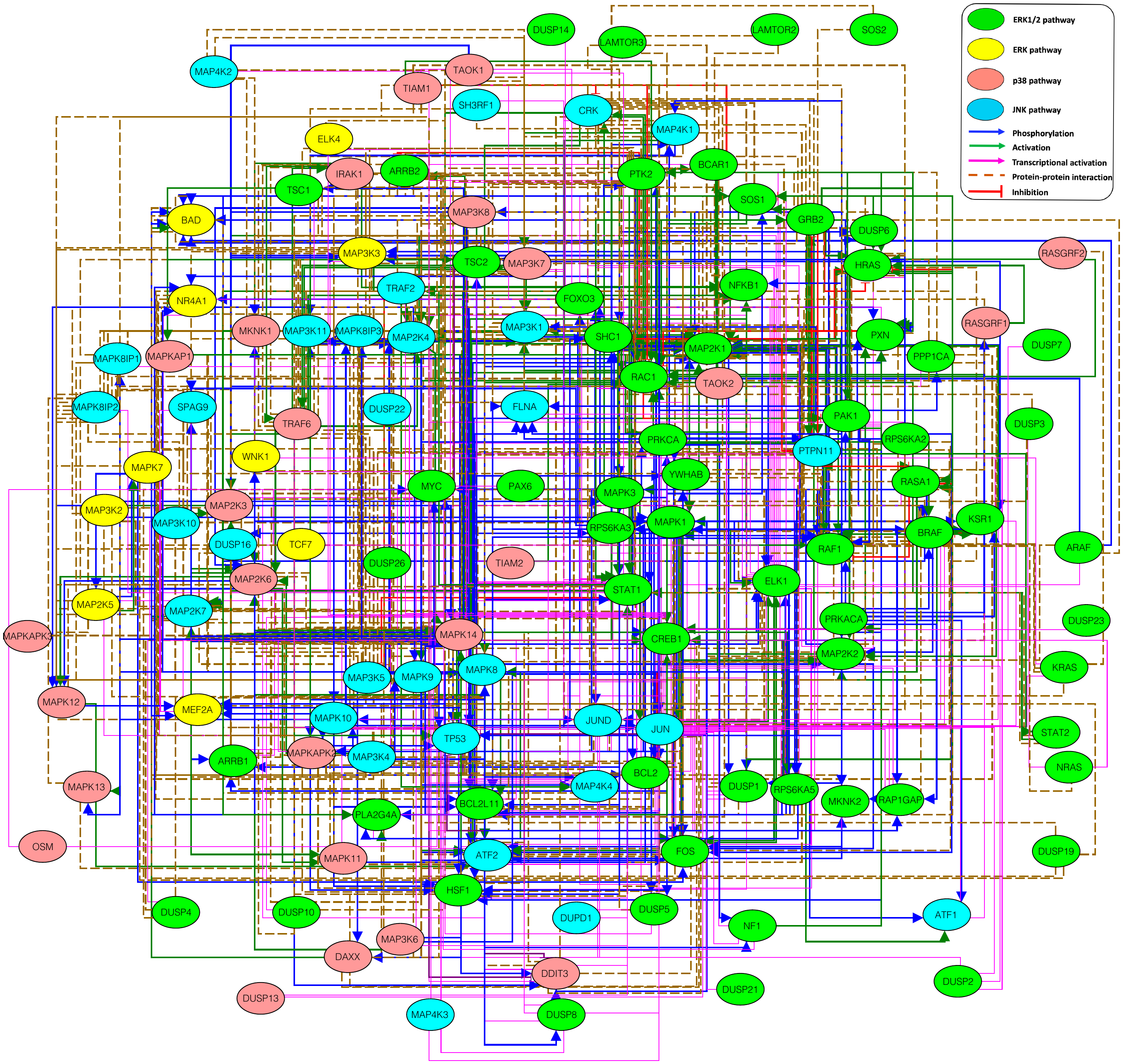

**Supplementary Figure 1:** Complete connectivity of MAPK signalling pathways components. The nodes are coloured according to the MAPK pathway module (ERK1/2, p38, JNK or ERK5 pathways) in which the protein participates. The connectivity of network components was extracted from the BioGrid database, Kinase Enrichment Analysis database, Chromatin Enrichment Analysis database, and UCSC super pathway databases.

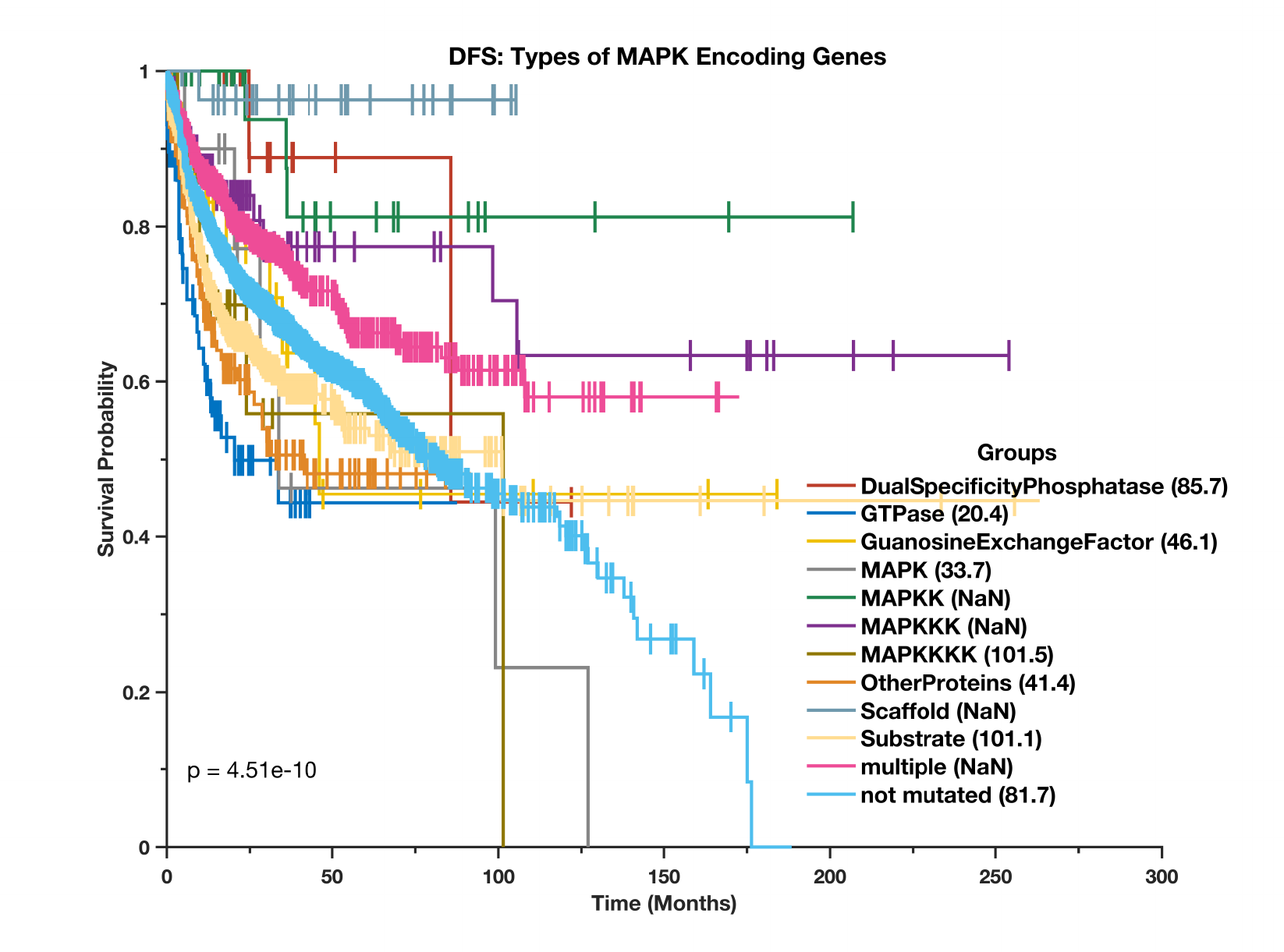

**Supplementary Figure 2:** Kaplan-Meier curve of the disease-free survival periods of patients with tumours that have mutations to genes that encode the various classes of MAPK proteins.

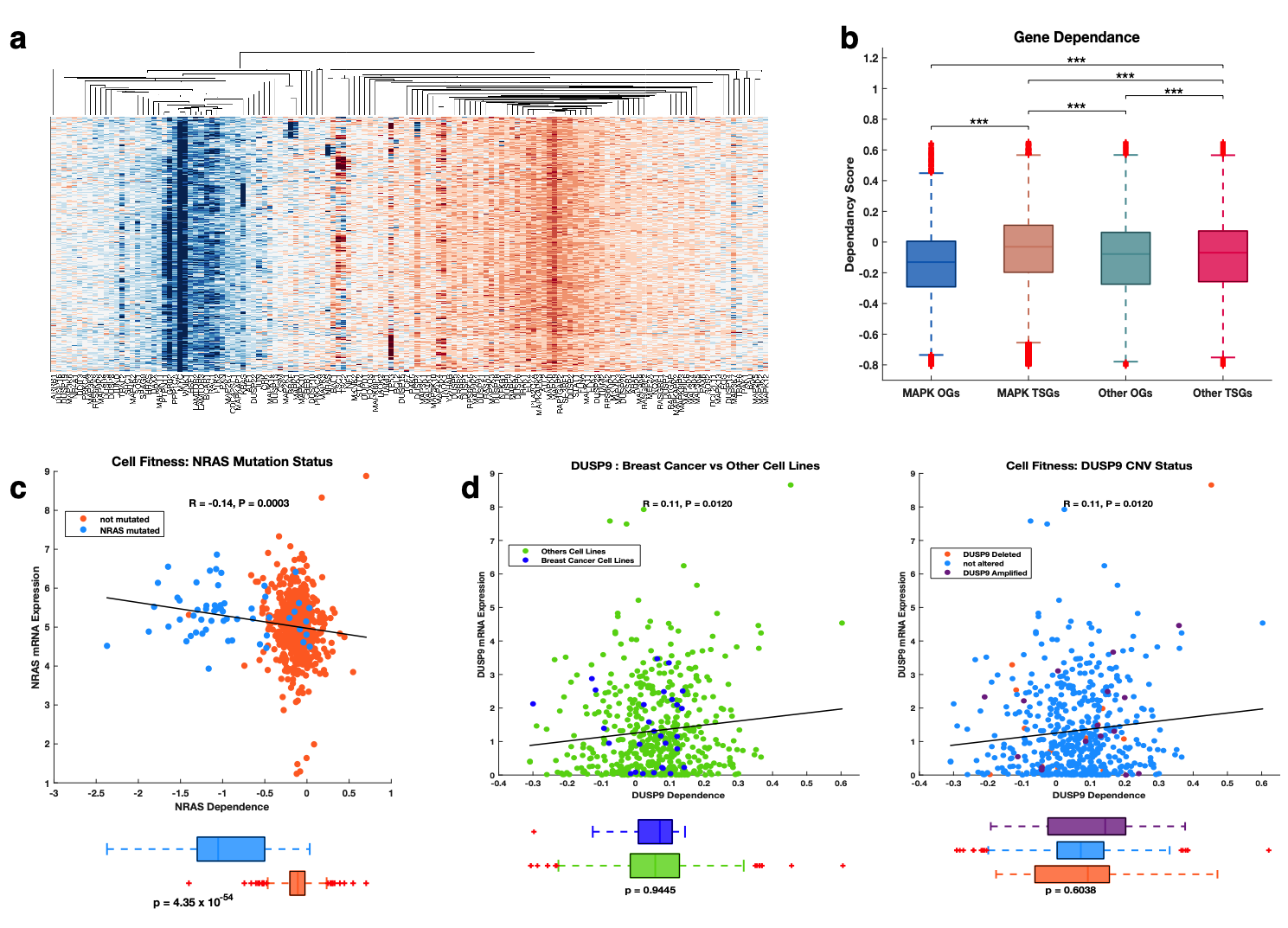

**Supplementary Figure 3: (a)** Clustergram of CRISPR-derived gene dependence scores across the cell lines. The bluer colour indicates reduced cell fitness and the redder colour indicates increased cell fitness after CRISPR gene knockout. The clustergram was produced using unsupervised hierarchical clustering with the Euclidean distance metric and complete linkage. **(b)** Mean difference in the CRISPR-derived gene dependence scores between four categories of genes: (1) MAPK pathway genes that are oncogenes, (2) MAPK pathway genes that are tumour suppressor genes, (3) oncogenes that are not MAPK pathway genes, and (4) tumour suppressor genes that are not MAPK pathway genes. The CRISPR-derived gene dependence scores were compared between these groups using one-way analysis of variance. *** denote statistical significance for adjusted p-values for each pairwise comparison using the Student’s *t*-test. On each box, the central mark indicates the median, and the bottom and top edges of the box indicate the 25th and 75th percentiles, respectively. The whiskers extend to the most extreme data points not considered outliers, and we plotted the outliers individually using the '+' symbol. **(c)** Correlation and the mean difference in *NRAS* dependence scores between cell lines that harbour *NRAS* mutations and those that do not harbour *NRAS* mutations. **(f)** Correlation between *DUSP9* mRNA transcript levels and *DUSP9* dependence score, and the mean difference in *DUSP9* dependence score between breast cancer (the cancer type with most *DUSP9* mutations) cell lines and all other cancer cell lines. **(g)** Correlation between *DUSP9* mRNA transcript levels and *DUSP9* dependence score and the mean difference in *DUSP9* dependence between the cell lines that gained copies of the *DUSP9* gene, have lost copy of the *DUSP9 gene*, and these with normal copies of the *DUSP9* gene.

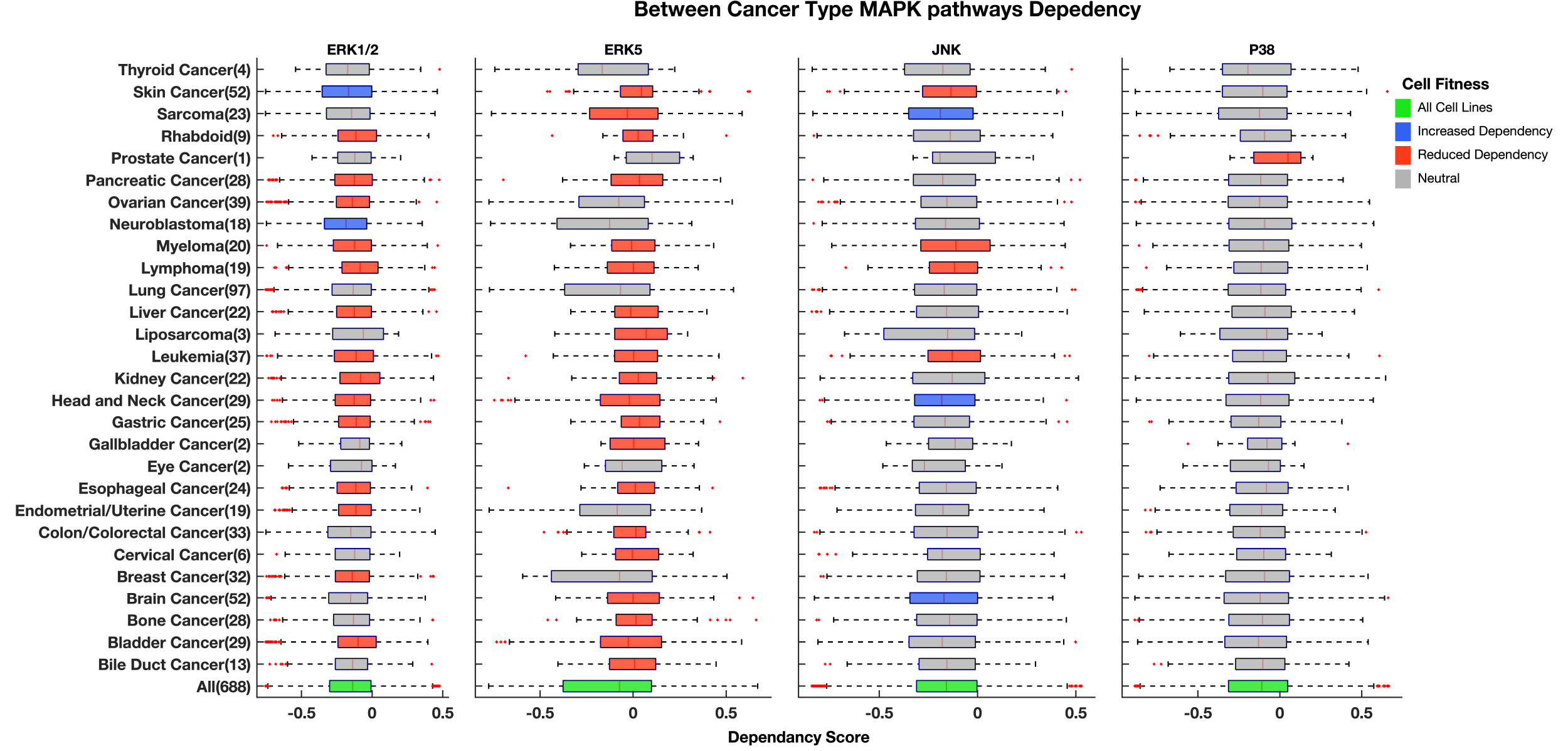

**Supplementary Figure 4:** Mean difference comparisons between the pooled CRISPR-derived gene dependence scores of each cancer types for a particular MAPK signalling pathway module vs the pooled CRISPR-derived gene dependence scores of all the genes that are not MAPK pathway genes for all 688 of the cancer cell lines (the green coloured boxplots). The blue boxplots indicate statistically significant loss of cell fitness and the red boxplots statistically significant increased cell fitness. P-values for each comparison were calculated from Welch's t-test. On each box, the central mark indicates the median, and the left and right edges of the box indicate the 25th and 75th percentiles, respectively. The whiskers extend to the most extreme data points not considered outliers, and the outliers are plotted individually using the '+' symbol.

**Supplementary Figure 5:**  Mean difference comparisons between **(a)** the pooled CRISPR-derived gene dependence scores of each cancer type for genes that encode the classes of MAPK pathway protein versus the pooled Achilles dependence score for those genes across all 688 of the cancer cell lines (the green coloured boxplots). **(b)** The pooled CRISPR-derived gene dependence scores of each cancer type for genes that encode the classes of MAPK pathway protein versus the pooled CRISPR-derived gene dependence scores of all the genes that are not MAPK pathway genes across all the 688 cell lines. Note that we made all comparisons with the pooled dependence score across all the 688 cancer cell lines (the green coloured boxplots). See Supplementary Figure 5 for description of the boxplots.

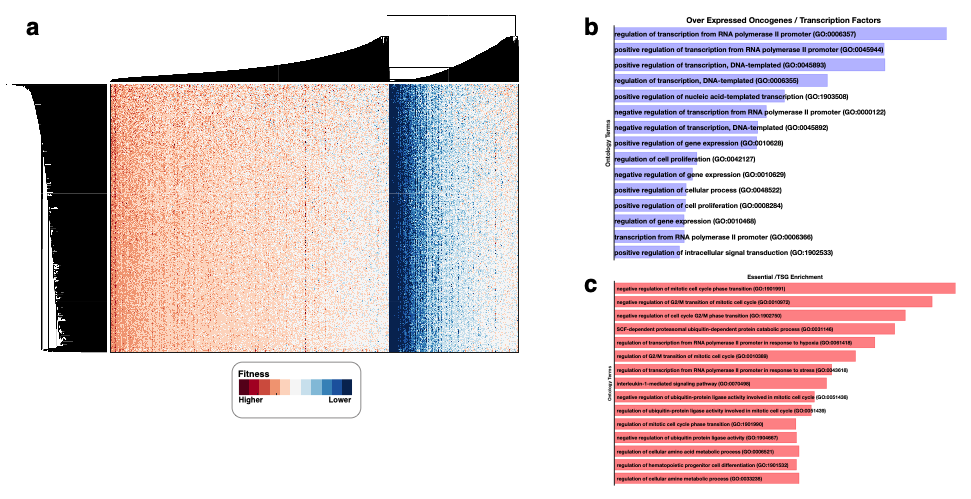

**Supplementary Figure 6: (a)** Clustering of the CRISPR-derived gene dependence scores across the 18023 genes that were profiled by the Achilles project. The bluer colour indicates a reduced fitness, whereas the redder colours indicated increased fitness of the cell lines after CRISPR genes knockouts. **(b)** Showing the top-ranked gene ontology biological process terms enriched for gene sets that showed a negative linear Pearson's correlation coefficient < -0.3. The genes list is composed of mostly oncogenes and transcription factors such as the *MYC* proto-oncogene (see Figure 5b). Also, see Supplementary File 3 for the complete list of gene ontology biological process terms that we found enriched. **(d)** Showing the top-ranked gene ontology biological process terms enriched for the gene sets that showed a positive linear Pearson’s correlation coefficient > 0.3. The gene list is composed of mostly tumour suppressor genes such as *CDKN2A*. Also, see Supplementary File 3 for the complete list of gene ontology biological process terms that we found enriched.

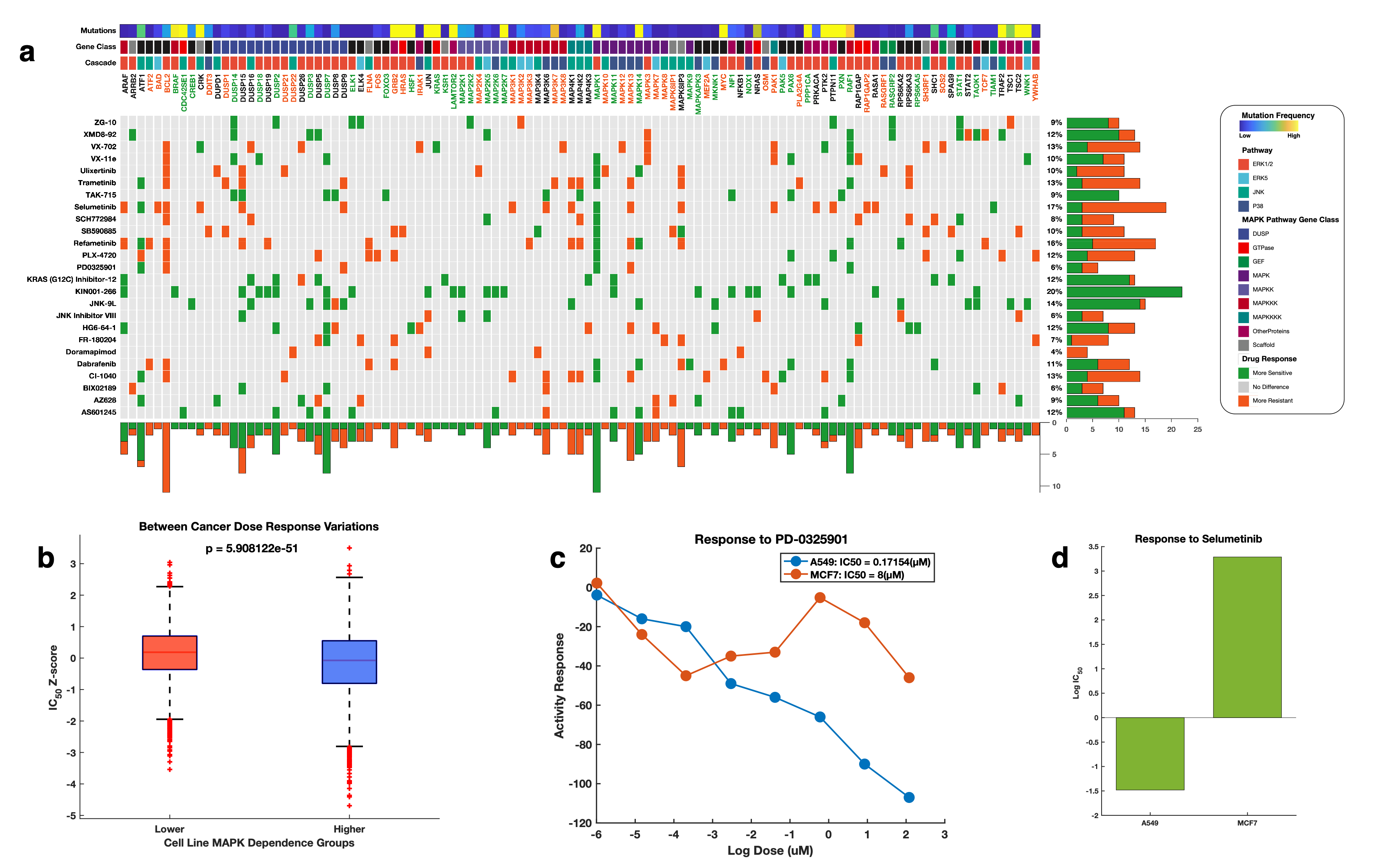

**Supplementary Figure 7: (a)** The integrated plot is showing the relationship between gene mutations and the drug response of the cancer cell lines. From top to bottom, the panel indicate: Mutations; the overall mutations frequencies observed for the gene along that column. Gene Class; the class of the MAPK pathway protein encoded by the gene. Pathway Module; the MAPK pathway module in which the gene participates. Heatmap; The marks on the heatmap are coloured based on how mutations in the gene in each column affect the efficacy of the drug given along each row: (1) with green denoting significantly increased sensitivity between cell lines that have mutations and those that do not have mutations in the gene, (2) grey for no statistically significant difference, and (3) orange denoting significantly increased resistance. The gene (column labels) are coloured based on the overall calculated effect of mutations on the efficacy of the drug given along each row: green; all the cell lines are significantly more sensitive to all the MAPK pathway inhibitors, orange; all the cell lines are significantly more resistant to all the MAPK pathway inhibitors, and black; a mixed response to MAPK pathway inhibitors. The bar graphs represent the total numbers of drugs whose dose-responses are significantly increased (green) or decreased (orange). **(b)** Overall comparison of the dose-responses to MAPK inhibitors between the cell lines with higher dependence on MAPK signalling and those with lower dependence on MAPK signalling. **(c)** Dose-response of the A549 and MCF7 cell lines to PD-0325901 as profiled by the GDSC. (d) Dose-response of the A549 and MCF7 cell lines to selumetinib as profiled by the CCLE.
